# Interaction of dengue NS3 with human RNA silencing machinery through HSPA1A

**DOI:** 10.1101/2020.06.08.140590

**Authors:** Pavan Kumar Kakumani, Rajgokul K. Shanmugam, Mahendran Chinnappan, Inderjeet Kaur, Arun P. Chopra, Pawan Malhotra, Sunil K. Mukherjee, Raj K. Bhatnagar

## Abstract

Viruses encode multiple proteins that interact with different host factors to aid in their establishment inside the host. Viral Suppressor of RNA silencing (VSR) are one such class of proteins that have been shown to interact with components of host machinery involved in post transcriptional gene silencing, a known antiviral defence mechanism. In the present study, we showed that dengue NS3, a known VSR not only interacts with HSPA1A, a cellular chaperone, but also modulates its expression levels. Further, we revealed HSPA1A associated with host RNA silencing machinery through its interaction with Argonaute proteins; Ago1, Ago2 and co-localizes with them in the cytoplasm of the cell. Together, these results provide evidence for involvement of other host partners in mediating VSR function of dengue NS3 and aid in deeper understanding of mechanisms underlying viral suppression of RNA silencing.

## Introduction

Infection and pathogenesis of viruses are related to host response that gets activated within a cell. Host RNA silencing is one of many interconnected pathways that is activated upon viral infection [1–5]. Most viruses counter the RNAi pathway by expressing the proteins referred as viral suppressors of RNA silencing (VSRs) that interfere with RNAi pathway [6,7]. VSRs have been well characterized in many plant and invertebrate viruses, however, their existence/role is a matter of contention among mammalian viruses, despite the fact that all components of RNAi machinery necessary for anti viral defence are conserved in mammals [4]. Role of RNAi as an anti viral mechanism in mammalian cells has been strengthened by a number of recent studies. A detailed analysis of virus derived small RNAs and host derived short RNAs in Hepatitis C, Polio, Dengue, Vesicular stomatitis and West Nile viruses provided convincing evidence for the role of RNAi in viral pathogenesis and host response [8]. Additionally, a number of reports have also shown a correlation between the levels of RNAi factors and virus infection both in mammalian cell lines and in animal models [5].

VSRs are reported to be multi-functional proteins that act at different levels to evade the host silencing pathways and to promote viral replication. Many VSRs have been identified from different animal viruses, but, a few like Ebola virus and HIV1 encode multiple VSRs. The VSRs from Ebola virus; VP30, VP40 and VP45 interact with multiple host RNAi proteins to evade the silencing response, while Tat and Rev proteins from HIV1 suppress both the Dicer and Ago dependent silencing pathways [4, 6, 9–12]. Recently, we have reported the presence of multiple suppressors from Dengue virus, namely, NS4B and NS3. Dengue NS4B, an IFN antagonist was shown to possess VSR properties, which interferes with dicing activity of processing long dsRNA substrates into siRNAs both *in vitro* and in cell line based assays [13]. NS3, a serine protease-helicase, was also shown to suppress the RNA silencing machinery by interfering with Ago1 loading of miRNAs [14]. Besides our recent report on interaction of dengue virus NS3 with the molecular chaperone, HSC70 to facilitate conditions favourable for viral replication, a number of reports have already shown that, surface heat shock proteins such as, HSP90 and HSP70 are part of Dengue virus receptor complex in human cells [15]. Heat shock treatment of monocytic U937 cells results in increased infectivity of Dengue virus in these cells and HSP70 positively regulates dengue replication in THP1 cell lines [16, 17].

In view of these findings, in the present study, we explored for other interacting partners of NS3 in human cell lines. Immunoprecipitation followed by mass spectrometric (MS) analysis in dengue NS3 expressing HEK293T cells identified HSPA1A, a host chaperone, as an interacting partner of NS3. As HSC70/HSP90 complex has been shown to mediate ATP-dependent RISC loading and NS3 interferes with loading of miRNAs into Ago1 protein, we next analysed the association of HSPA1A with host RNA silencing machinery. These findings along with earlier reports thus provide insights into the mode of action NS3 as a VSR, mediating the host RNAi suppression mechanism.

## Materials and methods

### Co Immunoprecipitation

To identify the host interacting partners of dengue NS3, cells expressing HA-NS3 were lysed in IP Lysis/Wash buffer (0.025M Tris, 0.15M NaCl, 0.001M EDTA, 1% NP-40, 5% glycerol; pH 7.4 with 1X Halt Protease and Phosphatase Inhibitor Cocktail (Thermo Scientific, USA)) at 48 h post transfection and immunoprecipitated using HA tag antibody as per the instructions given in the Pierce Co-Immunoprecipitation kit (Thermo Scientific, USA).

To examine the interaction between dengue NS3 and the host factors, pull down assays were performed using HA-Tag Rabbit mAb (Sepharose® Bead Conjugate) (Cell Signalling Technology, USA). Briefly, 400μg of total protein from the cell lysate was incubated with the 20μl of Sepharose Bead Conjugate at 4°C over night under rotation. After the incubation, the pellet was washed five times with IP Lysis/Wash buffer and 30μl of 1X SDS loading buffer was added to heat at 95°C for 2-3 min. Later, the suspended pellet was centrifuged to extract the supernatant and loaded onto 10% SDS PAGE gel for immuno blotting.

### Sample preparation for MS analysis

The immune precipitated samples using HA antibodies were resolved by SDS-PAGE and the gel was silver stained using Pierce Silver Stain Kit (Thermo Scientific, USA). The distinct protein bands were excised and were subjected to in-gel trypsin digestion using In-Gel Tryptic Digestion Kit (Thermo Scientific, USA). The recovered peptides in digestion buffer were dried under vacuum and dissolved in 0.1% Trifluoroacetic acid (TFA) in Milli-Q water for mass spectrometric (MS) analysis.

### Western blotting

The protein samples were run on 10% SDS PAGE gel and transferred onto a nitrocellulose membrane. The membrane was incubated in blocking solution (3% BSA in PBS buffer+0.05% Tween 20) for 1 hour with gentle shaking at 4°C over night. The blocking solution was replaced with primary antibody solution (1:1000 dilution in PBS buffer) and incubated for 1hour at room temperature with gentle shaking. Thereafter, the membrane was washed thrice with PBS buffer and 0.05% Tween-20 for 5 min each. After washing, HRP conjugated secondary antibodies solution (1:2000 dilution in PBS buffer) was added to the membrane and incubated for 1 hour at room temperature with constant shaking. After that, membrane was washed thrice with PBS buffer and 0.05% Tween-20 for 10 minutes each. The protein-antibody complex was developed by adding 2-3ml of SuperSignal West Pico Chemiluminiscent substrate (Thermo Fisher, USA).

## Results and Discussion

More than 50 VSRs have been identified up till now and they differ in primary sequence and structure [4–7]. Many of these VSRs are multifunctional proteins and play important roles in viral replication, coating, movement and pathogenesis besides suppressing host RNAi response [6, 9]. VSRs suppress RNA silencing via several mechanisms. Two most general mechanisms are through dsRNA binding to sequester small RNA duplexes and second one is through interaction with the components of RNA silencing pathways [18–20].

We have earlier shown that HSC70, a cellular chaperone, restricts dengue virus replication in human cell lines [14]. Heat Shock Proteins 90 & 70 were shown to be part of Dengue Receptor Complex and also participate in loading of small RNA duplexes into Ago proteins in both human cell lines [15, 21]. To investigate whether dengue NS3 interacts with other chaperone(s) besides HSC70, in the present study, we expressed dengue NS3 in HEK293T cells by transfecting a plasmid, NS3/pSELECT-NHA-zeo containing NS3 gene cloned downstream of HA-tag sequence. The cells transfected with empty plasmid, pSELECT-NHA-zeo served as vector control. Expression of NS3 was analysed by western blot analysis of transfected cell lysate using anti-HA antibody. As shown in Fig. 1B, HA-NS3 protein was expressed in cells transfected with NS3/pSELECT-NHA-zeo plasmid, while cells transfected with empty plasmid did not show expression of NS3. HEK293T cell expressing HA tagged NS3 and cells transfected with empty vector were lysed and the lysates were separately immunoprecipitated using anti-HA antibodies. Immunoprecipitates were resolved on SDS-PAGE gel and stained with colloidal blue to visualize the protein bands. Distinct bands were observed in HA-NS3 expressing cell lysate against the mock transfected at ~70 kDa size (Fig. 1A). These distinct bands were excised and in gel trypsin digested for mass spectrometric analysis. The mass analysis identified a protein HSC70, a known interacting partner of dengue NS3 along with another host protein, HSPA1A, also a molecular chaperone. The peptide sequences and % coverage of the protein used to annotate the host component, HSPA1A are given in Fig. 1C, D. To further confirm the interaction between dengue NS3 and HSPA1A, HEK293T cells were co-transfected with two plasmids, NS3/pSELECT-NHA-zeo and pcDNA5/FRT/TO V5 HSPA1A to co express HA-tagged NS3 and V5 tagged HSPA1A, respectively. Forty eight hours post transfection, the cells were lysed in non-denaturing conditions and co-immunoprecipitation was performed using HA-Tag Rabbit mAb Sepharose Bead Conjugate (Cell Signalling Technology Inc., USA). The elute and cell lysate were resolved on SDS-PAGE gel and western blotting was performed employing HA-tag and V5-Tag antibodies. As shown in Fig. 2A, V5-HSPA1A was pulled along with HA-NS3, but not in mock transfected cells. The expression of HA-NS3 and V5-HSPA1A was also confirmed in the cell lysate. These results clearly indicated that, dengue NS3 protein interacts with HSPA1A. Such an interaction between HSPA1A and NS3 from JEV, another flavivirus has been previously reported [22].

**Figure 1:**
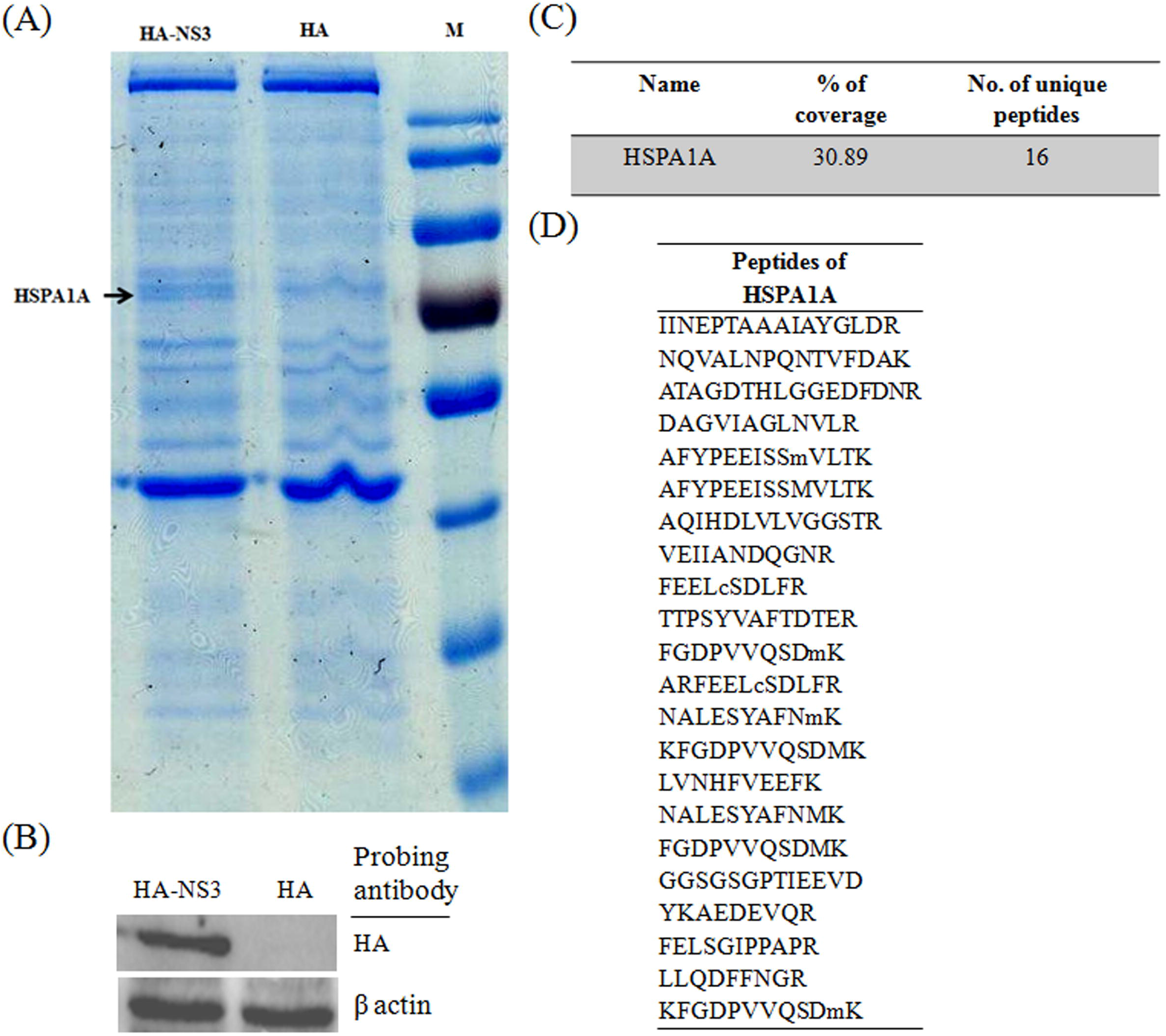
Identification of dengue NS3 host partners from GFP reversion assays: (A) SDS-PAGE gel stained in colloidal blue showing the distinct protein bands processed for MS analysis. (B) Western blot analysis of the pulled down samples and the cell lysate checking the expression of HA-NS3. (C, D) The tabular format, giving the details of % coverage and the peptide sequences used to annotate the host protein, HSPA1A.

**Figure 2:**
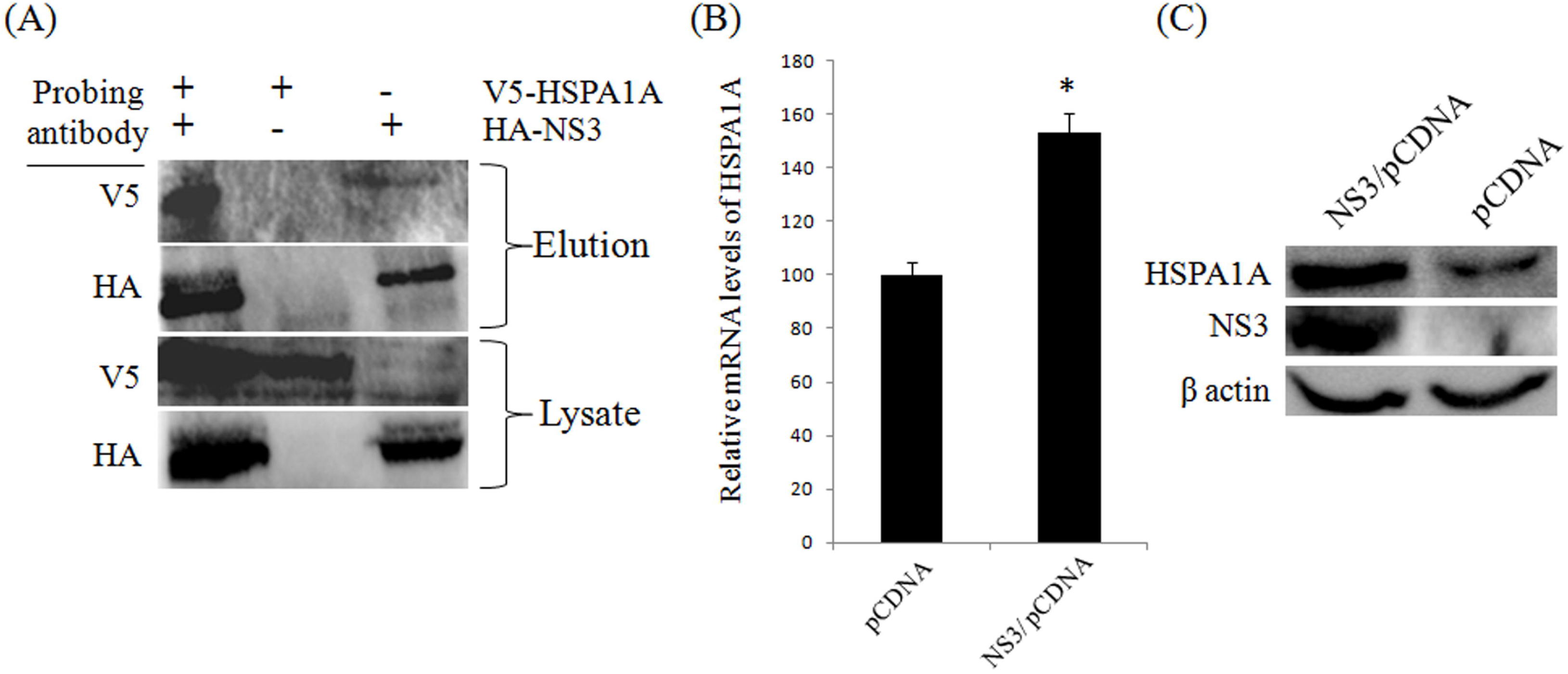
Association between dengue NS3 and HSPA1A: (A) Western blotting of immunoprecipitates from the HEK 293T cells over expression HA-NS3 and V5-HSPA1A (B) qRT-PCR analysis of relative mRNA levels of HSPA1A upon over expression of dengue NS3 in HEK293T cells. GAPDH served as endogenous control. The statistical significance of the values was determined by the (*) P value <0.05 (C) Western blotting of total protein from HEK293T cells over expressing dengue NS3.

Two independent reports have shown up regulation of HSPA1A upon dengue infection in human monocytic THP1 cells [17] and positive regulation in Japanese encephalitis virus (JEV) replication [22]. To know whether the up regulation of HSPA1A is due to over expression of dengue NS3, we examined the expression levels of HSPA1A upon over expression of dengue NS3 in HEK293T cells. Briefly, HEK293T cells were transfected with NS3/pSELECT-NHA-zeo and another set were mock transfected with the empty vector, pSELECT-NHA-zeo. Forty eight hour post transfection, the cells were lysed and both the protein and total RNA were isolated separately. The total RNA treated with DNase I was processed for qRT-PCR to determine the mRNA levels of HSPA1A. As shown in Fig. 2B, we observed ~ 56% increase in its mRNA levels relative to GAPDH. Further, westerns confirmed the enhanced protein levels of HSPA1A (Fig. 2C) in NS3 expressing HEK293T cell lines. These results were further corroborated by careful analysis of the microarray data of mRNAs in dengue NS3 expressing HEK293T cells as reported earlier [14]. Table S1 shows alterations in molecular chaperone levels upon expression of dengue NS3 in HEK293T cells, suggesting an interplay between dengue NS3 and HSPA1A. Together these results indicate that, during dengue viral infection when dengue NS3 is expressed, there is an increase in the expression of HSPAIA that might facilitate a better interaction between both the proteins for an affective suppression mechanism at Ago level. However, further studies are warranted to fully understand the significance of this interaction in the physiological context of virus infected human host.

We next looked into the relation if any between HSPA1A and the host RNA silencing machinery. Heat shock proteins like HSC70/HSP90, Fkbp4/5 were shown to be involved in the loading of miRNAs into Ago proteins and the facilitation of RISC complex respectively [21, 23]. To elucidate the association of heat shock proteins with functional RNAi protein complexes; we performed pull down assays by over expressing HA-Ago1 and HA-Ago2 in HEK293T cells. Briefly, the cells were transfected with plasmids, pIRESneo-FLAG/HA Ago1 or pIRESneo-FLAG/HA Ago2 or with empty vector, pIRESneo-FLAG/HA. Forty eight hour post transfection, the transfected cells were lysed in non-denaturing conditions and pull down assays were performed using HA-Tag Rabbit mAb Sepharsoe Bead Conjugate. The immunoprecipitates were resolved on SDS PAGE gel and silver stained to visualize the protein bands. As shown in Fig. 3A, distinct bands were observed on SDS-PAGE gel in Ago1 and Ago2 over expressing lanes in comparison to mock transfected cells. These distinct bands were excised and in gel trypsin digested for mass spectrometric analysis. The analysis identified both the Ago proteins along with HSC70, a known Ago associated host chaperone [21] along with HSPA1A. The peptide sequences and % coverage of the proteins used to annotate the hitherto unknown Ago associated factor, HSPA1A is presented in Fig. 3B, C.

**Figure 3:**
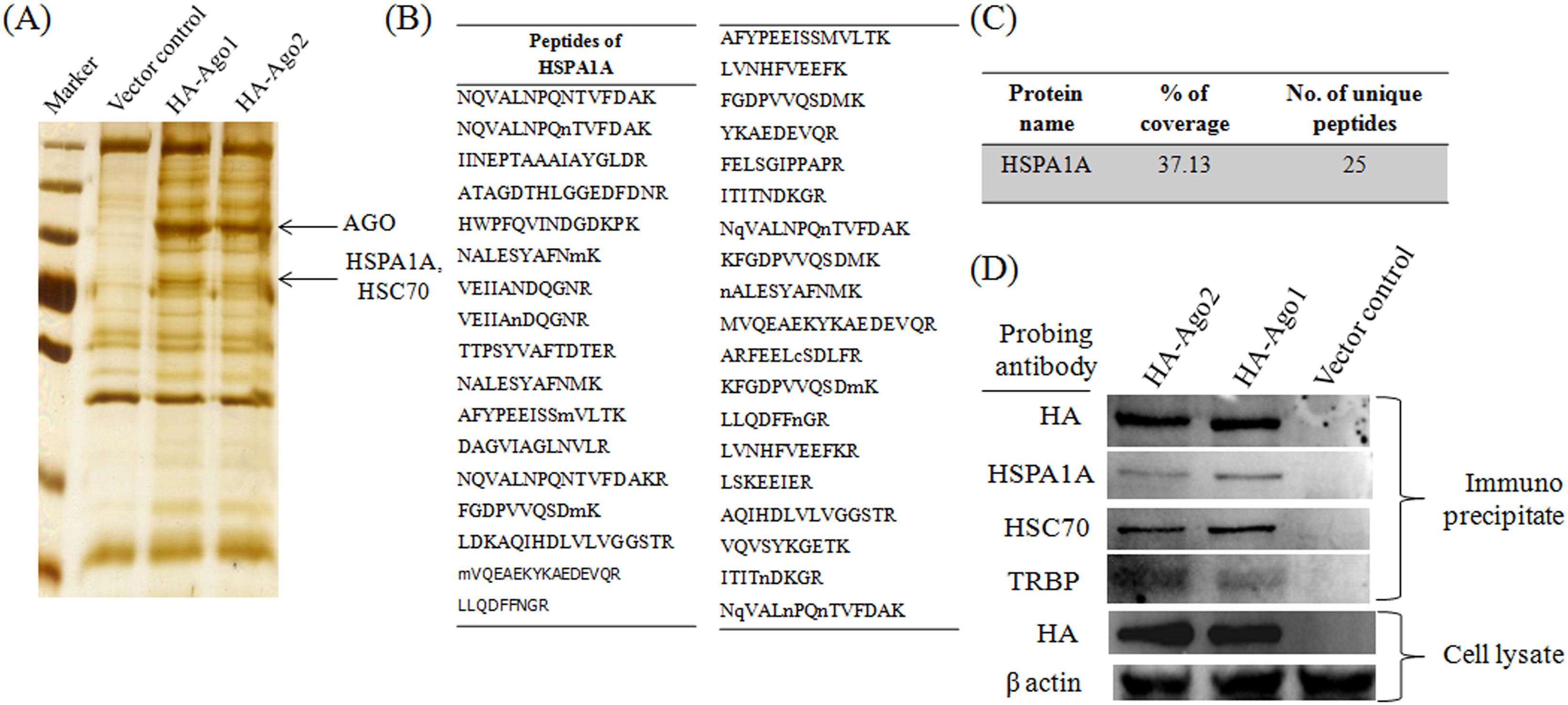
Association of HSPA1A with host RNA silencing machinery: (A) SDS-PAGE gel stained in silver stain showing the distinct bands processed for MS analysis. (B, C) The tabular format, giving the details of % coverage and the peptide sequences used to annotate the host protein, HSPA1A. (D) Western blotting of immunoprecipitates from HEK293T cells over expressing HA-Ago1 and HA-Ago2.

The peptide sequences and % coverage of Ago proteins and their associated factor, HSC70 is given in Table S2. To further confirm the results of MS analysis, the immunoprecipitates were subjected to western blotting using respective antibodies. As shown in Fig. 3D, HSPA1A was pulled down along with both the Ago proteins and the associated factors, HSC70 and TRBP. To further confirm the association HSPA1A with RISC components, we performed immunohistochemical studies to examine the localization of HSPA1A with Ago1 or Ago2. Respective antibodies against Ago1, Ago2 and HSPA1A were used to localize them, while the cell nucleus was highlighted in blue with DAPI. As shown in Fig. 4, HSPA1A was stained with red, while both the Ago proteins stained with green and the merged, highlighted in yellow revealed the co-localization of HSPA1A with both Ago1 and Ago2 mostly in the cytoplasm. These findings thus revealed that, HSPA1A is associated with Argonaute proteins and it would be interesting to investigate the role of HSPA1A in the loading of miRNAs as it was known to interact with HSC70 [24, 25]. Since, dengue NS3 was shown to affect the loading of miRNAs through its interaction HSC70, it would be of greater importance to elucidate the HSC70-NS3-HSPA1A association in regulating the miRNA loading for an effective repression of dengue viral host factors to establish dengue pathogenesis.

**Figure 4:**
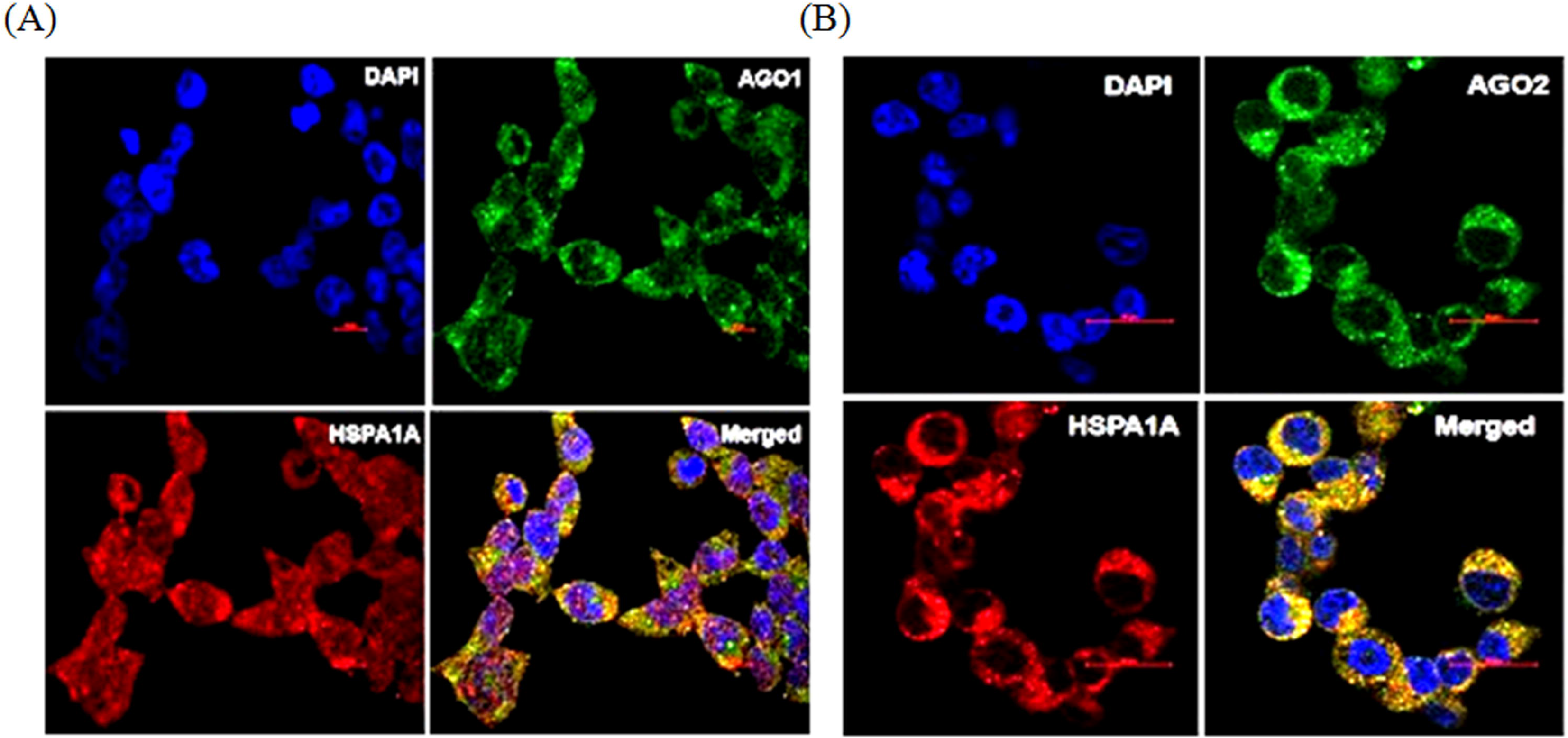
Co localization of HSPA1A with Ago1 and Ago2: (A, B) Sub cellular localization of Ago1, Ago2 (Green) and HSPA1A (Red) in HEK293T cells, as determined by anti Ago1, Ago2 and anti HSPA1A immunofluorescence microscopy. The co localization of HADHA with Ago1, Ago2 shown in merged (Yellow).

Present study thus identifies a new auxiliary factor of RNAi pathway, HSPA1A, a chaperone through which Dengue virus probably manipulates the host RNAi response. The results further show that dengue virus manipulation of RNAi pathway occurs through NS3, a viral suppressor protein. Overall, the results highlight the multiple interactions of dengue virus suppressor protein, NS3 with a number of host factors of RNAi pathway. It is intriguing to know, how the viral proteins associate themselves to the auxiliary factors rather the core catalytic components to suppress the host response. It may be because of their in ability to overcome the stronger interaction/functional compatibility between the core and the auxiliary components to yield an effective response from the host point of view. With the present findings corroborating our previous reports involving a larger interactome of host partners with viral proteins, it would be interesting to study systemic facilitation of these molecules by the viral components to the advantage of viral replication. Also, it may aid in identifying any new factors in understanding the host RNA silencing pathways as well as their conditional over expression and facilitation under stress conditions.

## Abbreviations

HSPA1A: heat shock 70KDa protein 1A
NS3: non-structural protein 3
miRNA: microRNA
siRNA: small interfering RNA
VSR: Viral Suppressor of RNA silencing
Ago: Argonaute
TRBP: TAR RNA binding protein
RISC: RNA induced silencing complex
HEK: human embryonic kidney
MS: mass spectrometric
VSR: viral suppressor of RNA silencing
IFN: interferon

## Acknowledgements

We thank Ms. Purnima Kumar, ICGEB New Delhi for her help with confocal microscopy during immunohistochemical analysis. We also thank Dr. Thomas Tuschl, The Rockefeller University, USA for providing the plasmids, pIRESneo-FLAG/HA Ago1 and pIRESneo-FLAG/HA Ago2 through Addgene Inc., USA. This study was funded by a financial grant from Dept. of Biotechnology, Govt. of India (BT/PR10673/AGR/36/579/2008).

## Author contributions

PKK, PM, SKM, RKB conceived the study designed the experiments. PKK and RKS performed the experiments. MC performed immunohistochemical studies. IK carried out the mass spectrometric analysis. APC provided guidance during the experiments and assisted in writing the manuscript. PKK, PM, SKM, RKB wrote the manuscript.

## Table Captions

**Table S1:** List of various chaperons altered upon NS3 expression in HEK293T cells.

**Table S2:** Peptide sequences and % coverage used to annotate HSC70 in mass spectrometric analysis.

