## Supplemental Tables for "Interaction of dengue NS3 with human RNA silencing machinery through HSPA1A"

**Table S1**

| **Fold Change**  **(NS3 vs Control)** | **Regulation** | **Gene Symbol** | **P value** |
| --- | --- | --- | --- |
| 1.206608 | up | HSBP1 | 0.020561 |
| 1.489153 | up | HSBP1L1 | 0.00576 |
| 1.865832 | up | HSPA1A | 0.007684 |
| 57.9581 | up | HSPA6/HSPA7 | 6.48E-04 |
| 1.551031 | up | HSPB1 | 0.029943 |
| 1.229522 | down | HSPB8 | 0.029538 |
| 1.23638 | down | HSPA8 | 0.012862 |

**Table S2**

| **Protein** | **Peptide sequences** | **% Coverage** | **No. of Unique peptides** |
| --- | --- | --- | --- |
| HSC70 | NQVAmNPTnTVFDAK | 53.72 | 26 |
|  | TVTNAVVTVPAYFnDSQR |  |  |
|  | QTQTFTTYSDNQPGVLIqVYEGER |  |  |
|  | NQVAMNPTNTVFDAK |  |  |
|  | IINEPTAAAIAYGLDKK |  |  |
|  | NQVAmNPTNTVFDAK |  |  |
|  | IINEPTAAAIAYGLDK |  |  |
|  | NQVAMNPTNTVFDAKR |  |  |
|  | SFYPEEVSSMVLTK |  |  |
|  | QTQTFTTYSDNQPGVLIQVYEGER |  |  |
|  | NQVAmNPTnTVFDAKR |  |  |
|  | FDDAVVQSDMK |  |  |
|  | ARFEELNADLFR |  |  |
|  | STAGDTHLGGEDFDNR |  |  |
|  | MKEIAEAYLGK |  |  |
|  | LDKSQIHDIVLVGGSTR |  |  |
|  | NQVAMNPTnTVFDAK |  |  |
|  | FDDAVVQSDmK |  |  |
|  | DAGTIAGLNVLR |  |  |
|  | STAGDTHLGGEDFDnR |  |  |
|  | MVQEAEKYKAEDEK |  |  |
|  | TVTNAVVTVPAYFNDSQR |  |  |
|  | LYqSAGGMPGGMPGGFPGGGAPPSGGASSGPTIEEVD |  |  |
|  | NQVAmNPTNTVFDAKR |  |  |
|  | SFYPEEVSSmVLTK |  |  |
|  | VEIIANDQGNR |  |  |
|  | VEIIAnDQGNR |  |  |
|  | TTPSYVAFTDTER |  |  |
|  | mKEIAEAYLGK |  |  |
|  | NSLESYAFNMK |  |  |
|  | cNEIINWLDK |  |  |
|  | NQTAEKEEFEHQQKELEK |  |  |
|  | mVNHFIAEFK |  |  |
|  | SQIHDIVLVGGSTR |  |  |
|  | HWPFMVVNDAGRPK |  |  |
|  | TVTnAVVTVPAYFnDSQR |  |  |
|  | DAGTIAGLnVLR |  |  |
|  | FEELNADLFR |  |  |
|  | MVNHFIAEFK |  |  |
|  | NQTAEKEEFEHQQK |  |  |
|  | NqVAMNPTnTVFDAK |  |  |
|  | NSLESYAFNmK |  |  |
|  | nSLESYAFNmK |  |  |
|  | RFDDAVVQSDMK |  |  |
|  | VQVEYKGETK |  |  |
|  | LLQDFFNGKELNK |  |  |
|  | ITITNDKGR |  |  |
|  | LYqSAGGmPGGMPGGFPGGGAPPSGGASSGPTIEEVD |  |  |
|  | NSLESYAFnMK |  |  |
|  | FEELnADLFR |  |  |
|  | LLQDFFnGK |  |  |
|  | NQTAEKEEFEHQqK |  |  |
|  | LLQDFFNGK |  |  |
|  | LLQDFFnGKELNK |  |  |
|  | EIAEAYLGK |  |  |
|  | IInEPTAAAIAYGLDK |  |  |
|  | ITITnDKGR |  |  |
|  | LSKEDIER |  |  |
|  | LYQSAGGMPGGMPGGFPGGGAPPSGGASSGPTIEEVD |  |  |
|  | RFDDAVVQSDmK |  |  |
